# Intestinal bacteria modulate the foraging behavior of the Oriental fruit fly *Bactrocera dorsalis* (Diptera: Tephritidae)

**DOI:** 10.1101/311910

**Authors:** Mazarin Akami, Awawing A. Andongma, Chen Zhengzhong, Jiang Nan, Kanjana Khaeso, Edouard Jurkevitch, Chang-Ying Niu, Boaz Yuval

## Abstract

The gut microbiome of insects directly or indirectly affects the metabolism, immune status, sensory perception and feeding behavior of its host. Here, we examine the hypothesis that in the Oriental fruit fly *(Bactrocera dorsalis*, Diptera: Tephritidae), the presence or absence of gut symbionts affects foraging behavior and nutrient ingestion. We offered protein starved flies, symbiotic or aposymbiotic, a choice between diets containing all amino acids or only the non-essential ones. The different diets were presented in a foraging arena as drops that varied in their size and density, creating an imbalanced foraging environment. Suppressing the microbiome resulted in significant changes of the foraging behavior of both male and female flies. Aposymbiotic flies responded faster to the diets offered in experimental arenas, spent more time feeding, ingested more drops of food, and were constrained to feed on time consuming patches (containing small drops of food), when these offered the full complement of amino acids. We discuss these results in the context of previous studies on the effect of the gut microbiome on host behavior, and suggest that these be extended to the life history dimension.

**Importance and significance of the study:** The gut bacteria of tephritid fruit flies provide nutritional benefits to their hosts, by making essential amino-acids readily available. Foraging for food is risky, as active flies are exposed to predators and incur a considerable investment of time and energy. Therefore, making beneficial compromises between the feeding time and nutrient ingestion is a question of survival for the flies. Our study demonstrates how gut bacteria drive this behavior by allowing symbiotic flies to forage optimally while acquiring essential nutrients. This finding adds a novel step to the nexus connecting the insect gut, its microbiome, the nervous system, chemoreception to individual patterns of foraging.

## 1. Introduction

Animals forage for nutritional resources in order to satisfy their requirements for growth and reproduction (1, 2). This behaviour is constrained by spatial and temporal factors (biotic and abiotic), and modulated to a large extent by each organisms experiential and metabolic state (3). Evidence from numerous studies suggests that insects (and other arthropods) are capable of tailoring their foraging activity and ingestion of nutrients in a manner that corresponds to their specific requirements (4) (and references therein).

In insects, responses to environmental stimuli are modulated by substrate specific chemoreceptors, whose sensitivity is modulated by the nutritional status of the individual (5–7). Thus, for example, in the vinegar fly *Drosophila melanogaster*, flies deprived of amino acids exhibited an enhanced response to amino acids missing from their diets (8). Similarly, tephritid fruit fly sensory responses and foraging activity are affected by nutritional status (9–11).

The microbiome resident in the gut of arthropod (and vertebrate) hosts adds another layer of complexity to the modulation of behaviour in general and foraging behaviour in particular (12–14). This effect has been demonstrated along the various steps of the nexus connecting the gut and its microbiome to behaviour, through metabolism, the immune and nervous systems, and sensory receptors. Thus, in *D. melanogaster*, the microbiota has multiple impacts on metabolism such as immune homeostasis, lipid and carbohydrate storage and vitamin sequestration (15–17). These effects are extended to responses to food and ultimately affect foraging activity. In *Tenebrio molitor* mealworms, individuals whose immune system was activated by a pathogen consumed significantly more proteinaceous food than healthy individuals (18). Conversely, stinkbug *(Megacopta punctatissima)* nymphs that acquire symbionts after hatching exhibit lower activity levels than symbiont free nymphs (19). Recently, Wong *et al.* (20) demonstrated that the microbiome of *D. melanogaster* influences the olfactory sensitivity and foraging behaviour of hosts in a manner that apparently benefits the bacteria specifically (21). Remarkably, there is evidence that the nutritional status of the host interacts with the microbiome to control foraging behavior. In *D. melanogaster*, the absence of specific amino acids will trigger specific appetites for the missing nutrient. However, the presence of bacteria (that presumably could provide the missing amino acid), overrides such preferences (22).

Tephritid fruit flies (Diptera: Tephritidae) harbour communities of bacteria dominated by species of Enterobacteriacae (23). These microbes have been shown to be involved in Nitrogen fixation (24, 25), reproductive success (26, 27), temporal host range expansion (28), protection from pathogens (29) and detoxification (30).

Adult tephritid flies require a mixed diet consisting of carbohydrate and protein, or at least protein precursors. These nutrients are acquired by active foraging during daylight hours. Sugars are acquired from nectar, honeydew and fruit juices, while nitrogenous compounds are sourced by feeding on bird feces, or in some cases bacteria on the phylloplane (31). The presence of gut bacteria in adult flies contributes to their nutrition, specifically by brokering intractable sources of Nitrogen into essential amino acids. Thus symbiotic olive flies *(Bactrocera oleae)*, were able to produce eggs when provided only with non-essential amino acids, while aposymbiotic flies were unable to do so (32, 33). Foraging for food is risky, as active flies are exposed to predators and incur a considerable investment of time and energy. Accordingly, in the present study we examine the hypothesis that in the Oriental fruit fly *(Bactrocera dorsalis*), the presence or absence of gut symbionts will affect foraging behavior and nutrient ingestion. We offered protein starved flies, symbiotic or aposymbiotic, a choice between diets containing all amino acids or only the non-essential ones. The different diets were presented in a foraging arena as drops that varied in their size and density, creating an imbalanced foraging environment. We predicted that symbiotic flies would consistently choose the diet that was most profitable in terms of foraging time. Conversely, we predicted that flies lacking symbionts would be constrained to forage on diets containing all amino acids, while incurring costs of increased exposure and foraging time.

## 2. Materials and Methods

### 2.1 The Study Organism and its Microbiome

The oriental fruitfly *Bactrocera dorsalis* (Diptera:Tephritidae:) is one of the most invasive, multivoltine and polyphagous members of the Tephritidae family. This fly, causes considerable loss of cultivated crops in most western and eastern parts of Asia and attacks over 350 hosts worldwide (24, 34). Studies based on Polymerase Chain Reaction-Denaturing Gradient Gel Electrophoresis (PCR-DGGE) fingerprinting and High throughput pyrosequencing of the 16S rRNA gene have highlighted the prevalence of microbial communities inhabiting the gut and reproductive organs of this insect, and which play critical roles in host physiology and behavior (24, 35–37). Pyrosequencing analysis of the *B. dorsalis* microbiome reveals 172 Operational Taxonomic Units assigned to 6 phyla (with Firmicutes as the most abundant in adult stages), 5 families, and 13 genera (24, 35). The predominant bacterial family in most of the previous studies was Enterobacteriaceae from which many cultivable species were identified, such as *Klebsiella oxytoca, Enterobacter cloacae, Listeria, Morganella, Moraxella proteus, Providencia rettgerii, Citrobacter freundii, Pseudomonas aeruginosa, Enterococcus phoeniculicola*, and *Lactobacillus lactis* (27, 38, 39).

### 2.2 Fly rearing and handling

*Bactrocera dorsalis* wild strain larvae were collected from infested fruits from the experimental orange orchard of Huazhong Agricultural University (30°4’N and 114°3’ E). The larvae were carefully removed after peeling the orange fruits and allowed to develop into wheat-bran based larval artificial diet. The third instar larvae were allowed to pupate in sterile sand under laboratory conditions and the resulting adults were kept under rearing since 2014 (24). At each generation, flies were reared as described by Nash and Chapman (40) with slight modifications. Briefly, 100 adults’ males and females were housed in 5L cages at equal proportions. These cages were maintained under controlled environment: 12:12 light-dark photoperiod; temperature 26±3°C, and 57±5% relative humidity. Water was provisioned *ad libitum* and the wheat bran based diet consisted of Tryptone (25 g/L), Yeast extract (90 g/L), Sucrose (120 g/L), Wheat bran (250 g/L), Agar powder (7.5 g/L), Methyl-p-hydroxybenzoate (4 g/L), Cholesterol (2.3 g/L), Choline chlorite (1.8 g/L), Ascorbic acid (5.5 g/L) and 1 L of distilled Water.

### 2.3 Symbiotic and Aposymbiotic flies

Symbiotic and aposymbiotic flies were produced from the laboratory established colony. Newly emerged 1 day old flies were sex-separated and divided into two groups of 30 flies each. The symbiotic groups consisted of flies (males and females) fed with Sugar diet for seven days using 9 cm petri dishes presented in cotton wool and the second groups were also fed with the same sugar meal in cotton wool inoculated with antibiotics (3mcg/mL Norfloxacin and 5mcg/mL Ceftazedime) from the 4^th^ treatment day (33). The antibiotics were selected after *in vitro* susceptibility test to 7 bacterial isolates and their *in vivo* capacity to significantly clear the gut of *B. dorsalis* within four feeding days (Table 1). All the flies were starved for 24h before experiments.

**Table 1.** Antibiotics susceptibility testing by disc diffusion of gut bacteria isolated from the oriental fruit fly *Bactrocera dorsalis*

| No. | Antimicrobial Agent | Disc potency (µg/piece) | *Enterococcus faecalis* | *Klebsiella oxytoca* | *Bacillus cereus* | *Enterobacter cloacae* | *Pantoea dispersa* | *Proteus mirabilis* | *Staphylococcus sciuri* |  |
| --- | --- | --- | --- | --- | --- | --- | --- | --- | --- | --- |
| 1 | Ampicillin | 10 | R | I | S | I | R | R | R |  |
| 2 | Ceftazidime | 30 | S | S | S | S | S | S | S | * |
| 3 | Chloramphenicol | 30 | S | S | I | R | S | I | I |  |
| 4 | Ciprofloxacin | 5 | I | S | S | S | R | S | S |  |
| 5 | Clindamycin | 2 | R | I | R | R | R | R | R |  |
| 6 | Erythromycin | 15 | R | R | R | R | R | R | R | *** |
| 7 | Gentamicin | 10 | S | S | R | S | S | S | S | ** |
| 8 | Kanamycin | 30 | S | S | R | S | S | S | S | ** |
| 9 | Minocyclin | 30 | S | S | R | R | I | I | R |  |
| 10 | Norfloxacin | 10 | S | S | S | S | S | S | S | * |
| 11 | Ofloxacin | 5 | S | S | R | S | S | S | S | * |
| 12 | Penicillins | 10 | I | R | R | R | R | R | R | *** |
| 13 | Piperacillin | 100 | S | S | S | I | R | S | R |  |
| 14 | PolymyxinB | 300 | S | S | R | S | S | S | S | ** |
| 15 | Tetracyclin | 30 | S | S | R | R | R | R | I | *** |

### 2.4 Preparation of experimental diets

Three different diets were prepared. A full diet (F) containing all amino acids (essential and non-essential), sucrose, and minerals, required for an optimal maintenance and reproductive development of adult flies; a non-essential amino acid diet (NE) containing exclusively non-essential amino acids, sucrose and minerals. A Sugar diet (Su) consisting only of 60% sucrose and minerals, was provided before the experiments. The diet ingredients and preparation procedures were done as described by Ben-Yosef *et al*. (33).

### 2.5 Experimental procedures

Following the seven day preparatory period during which flies were fed only sugar (as described above), an individual fly from each treatment was transferred to a 20 × 20 cm cage and allowed to acclimatize for 20 minutes before introducing a pair of petri dishes containing combinations of two different diet types (Full or NE) at different densities (Fig. 1). To create different foraging environments, 25 drops of 1 μL volume (very small so as to force the flies to seek out many drops in order to become satiated), and 5 drops of 5 μL volume were pipetted onto each dish. Each treatment of the six treatments was replicated 15 times and each replicate consisted of observing the protein starved individual male or female (symbiotic and aposymbiotic) for 1 hour. To motivate foraging behavior, all the flies were starved for 24 hours before experimental trials. Data on latency (time from diet exposure to the initial landing), the number of flies which landed, the choice of diet made, the number of drops consumed, and the time spent feeding were recorded.

**FIG. 1.**
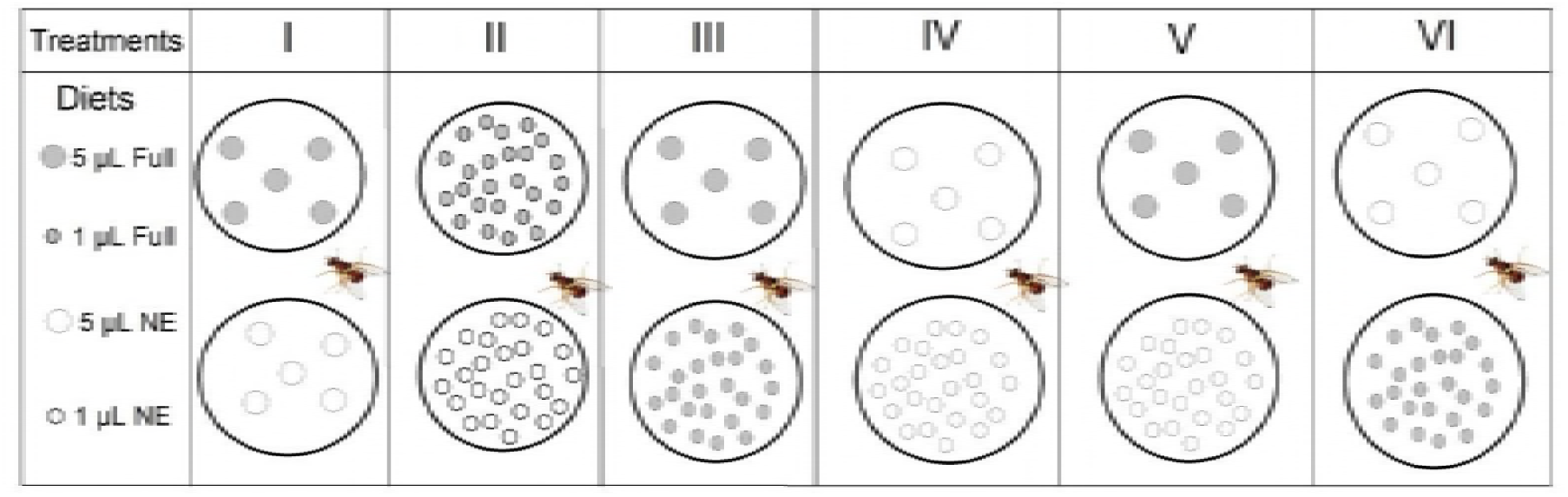
Experimental arenas of the effect of diet quality and density on foraging decisions by the oriental fruit fly *Bactrocera dorsalis.* Full diets contain all amino acids, and NE diets contain only the non-essential amino acids.

### 2.6 Statistical Analyses

All data were tested for homogeneity of variances using Levene’s tests. To determine the important factors that shape the foraging behavior of experimental flies, variables of overall response and latency were analyzed (each one separately) using the ordinary least squares regression model (SPSS 20.0 software) with sex, symbiotic status, treatment and the interaction between symbiotic status and treatment as effects. The One-Way Analysis of Variance (ANOVA) was performed to analyze data on landing, the number of drops consumed, and the time spent using SPSS 20.0 software (Statsoft Inc, Carey, J, USA). Duncan’s Multiple Range Test (DMRT) test was used for mean separations within and between each treatment. Differences among measured parameters were considered significant when the *p* values were less than 0.05 after comparison with DMRT. All results were expressed as the means with standard errors (SE), except data on the overall responses. OriginPro software version 8.5.1 was used to draw curves and graphs.

## 3. Results

### 3.1 Responses to the experimental arenas

The overall response (number of flies responding) to the different treatments was significantly affected by symbiotic status (Ordinary Least Squares Regression Model, F = 15.834; df = 3; r^2^ = 0.839; t = 6.048; P < 0.001), and by sex (F = 15.83; df = 3; r^2^ = 0.839; t = −2.946; P = 0.043) (Fig. 2A).

**FIG. 2.**
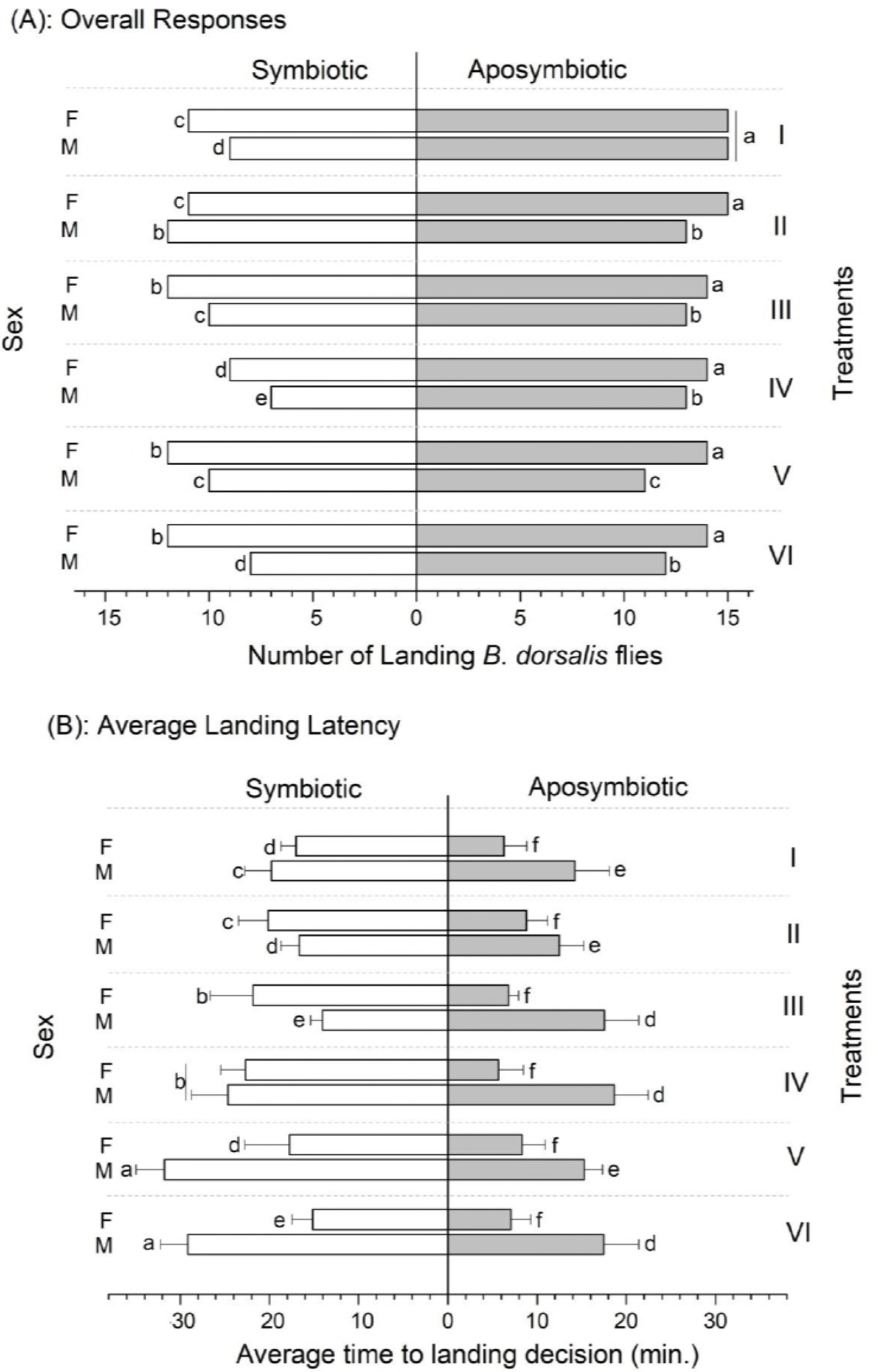
Response parameters of *Bactrocera dorsalis* to experimental diets as affected by the symbiotic status, sex and treatments. F: Females; M: Males. Means followed by different letters within and between responses or latencies are statistically different after Duncan’s Multiple Range Test at P = 0.05. Each response datum consisted of the overall number of flies which landed in different treatments regardless the diet quality or drop size. Each latency datum is presented as a Mean±SE of the overall latency of responded flies in each treatment. Each treatment consisted of 15 replications.

Furthermore, the latency to respond in the experimental chambers was similarly affected by symbiotic status (F = 11.538; df = 3; r^2^ = 0.796; t = −4.929; P < 0.0001) and sex (F = 11.538; df =3; r^2^ = 0.796; t = 3.24; P = 0.04) (Fig. 2 B). In general, aposymbiotic flies responded faster in the experimental chambers than the symbiotic ones, and females in aposymbiotic groups landed on food drops faster than the males (Fig. 2B).

The treatment itself did not affect the overall response of flies (F = 15.834; df = 3; r = 0.839; t = −1.498; P = 0.197), but significantly influenced the latency to respond (F = 11.834; df = 3; r^2^ = 0.796; t = 0.216; P < 0.001) (Fig. 2).

### 3.2 Switching behavior

Shifting from one diet to another was common, and showed a clear trend. No aposymbiotic flies which landed on the Full diet shifted to the NE diet and those which initially landed on the NE diet recorded a faster shifting latency (time to move from an initial patch to another) toward the Full diet (5.75±2.75 minutes, and 5.14±2.3 minutes in females and males, respectively) in comparison with the symbiotic females and males (15.75±2.25 minutes, and 12±0.75 minutes, respectively) (F = 4.54, t = 1.62; df = 14, P = 0.0016, and F = 2.65, t = −0.74, df = 12, P = 0.02, respectively, Table 2). However, 1 symbiotic female and male among those which initially landed on the Full diet shifted to the NE diet, but within a long shifting latency (28.25±3.5 minutes, F = 14.83, t = 2.8, df = 14, P = 0.001 and 19±4.2 minutes, F = 32.64, t = 1.79, P = 0.009, in females and males, respectively) (Table 2).

**Table 2.**
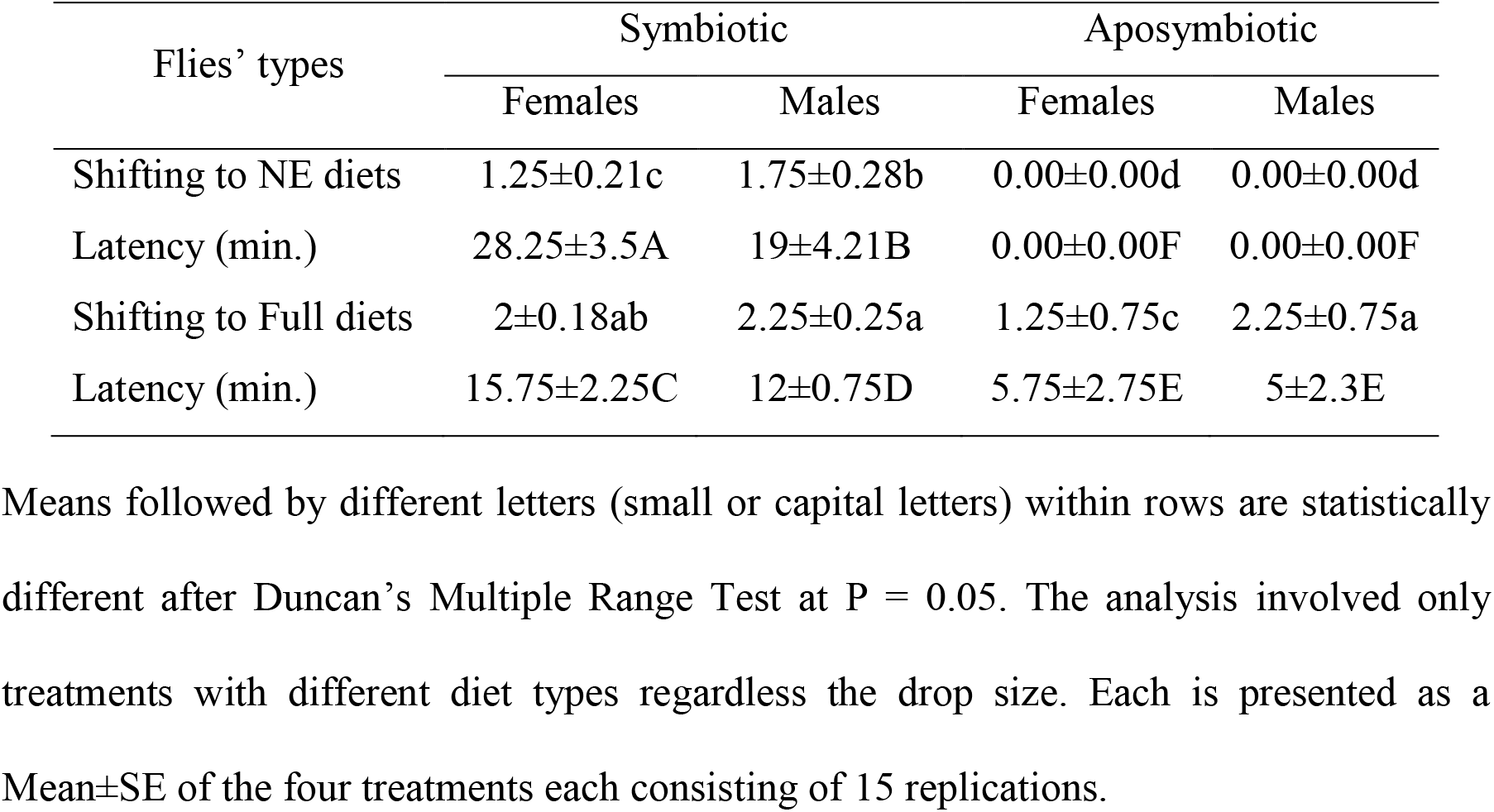
Switching behavior of *B. dorsalis* as influenced by diet quality.

| Flies' types | Symbiotic |  | Aposymbiotic |  |
| --- | --- | --- | --- | --- |
|  | Females | Males | Females | Males |
| Shifting to NE diets | 1.25±0.21c | 1.75±0.28b | 0.00±0.00d | 0.00±0.00d |
| Latency (min.) | 28.25±3.5A | 19±4.21B | 0.00±0.00F | 0.00±0.00F |
| Shifting to Full diets | 2±0.18ab | 2.25±0.25a | 1.25±0.75c | 2.25±0.75a |
| Latency (min.) | 15.75±2.25C | 12±0.75D | 5.75±2.75E | 5±2.3E |

### 3.3 Ingestion

Overall, the number of drops consumed depended significantly on drop size, diet (full or NE), treatment (foraging environment), sex and symbiotic status of the flies observed (ANOVA; F = 45.86, df = 5, P < 0.0001, r^2^ = 0.96).

Ingestion of Full diets (high or low volume) from all treatment groups was significantly higher in all tested flies, males and females (F = 64.12, df = 5, P < 0.0001, R^2^ = 0.94, and F = 11.72, df = 5, P < 0.0001, R^2^ = 0.83, respectively, Fig. 3 A-B). Nevertheless, compared to males, females displayed a significant preference toward diets with high reward (full diet, large drops) in unbalanced nutritional environments (F = 41.56, df = 5, P < 0.0001, r^2^ = 0.87, Fig. 3 A-B).

**FIG. 3.**
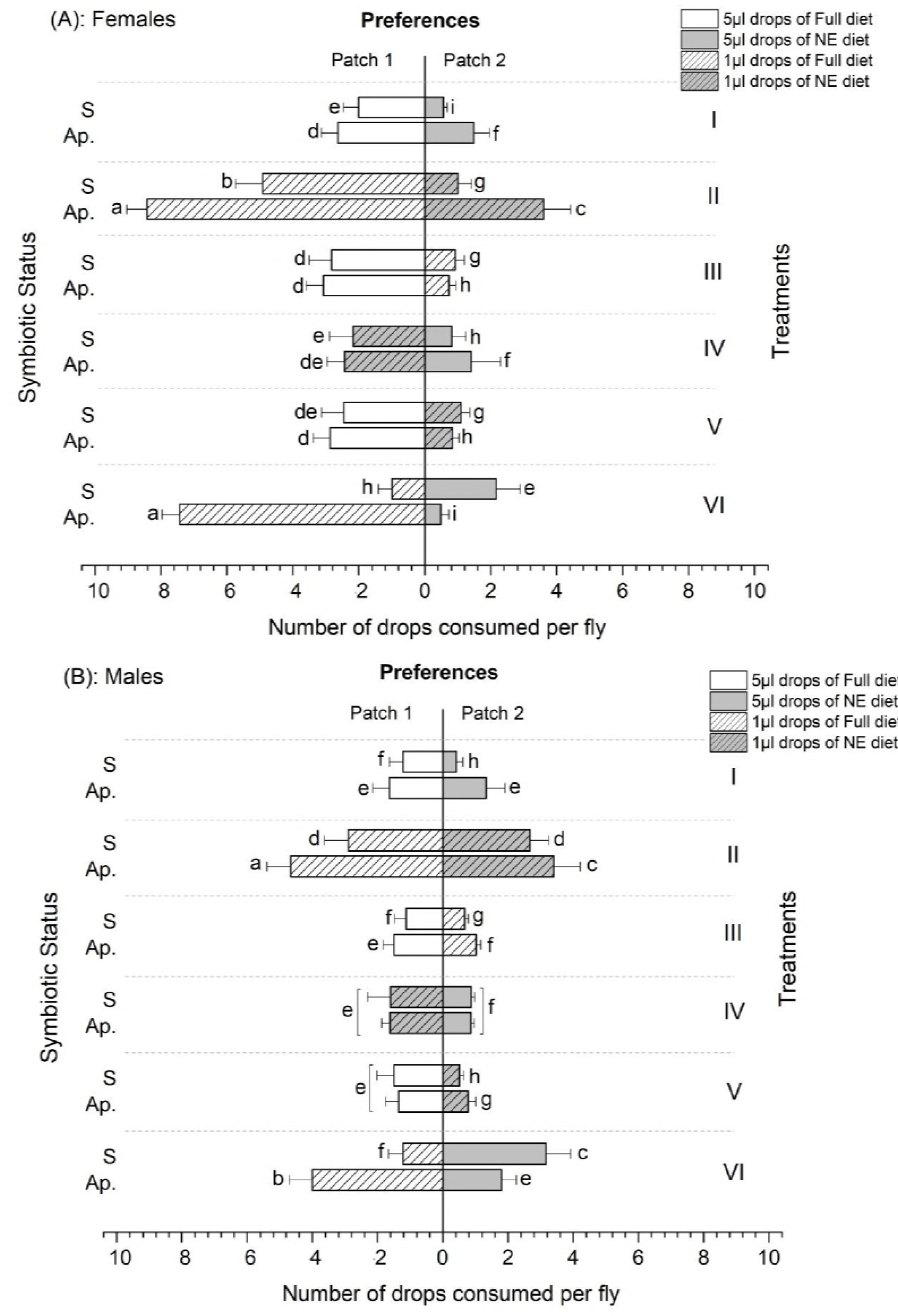
Number of nutritional drops consumed by symbiotic (S) and aposymbiotic (Ap.) *Bactrocera dorsalis* exposed to two diet patches (full and non-essential amino acids’ diets). Each bar represents the Mean±SE of consumed drops by each fly within an hour of observation. Mean values followed by different letters within and between treatments are statistically different after Duncan’s Multiple Range Test at P = 0.05.

Most importantly, aposymbiotic flies of both sexes preferentially chose to feed on the Full diet regardless of drop size. Thus, symbiotic condition significantly affected fly feeding behavior in treatment VI. Here, flies were offered many low volume drops of the Full diet, together with few high volume drops of the NE diet. Both male and female aposymbiotic flies were compelled to consume numerous drops of the low volume, Full diet drops. Conversely, Symbiotic flies of both sexes ignored the time consuming Full diet drops and consumed the larger drops containing the NE diet (F = 14.22, df = 5, P < 0.0001, R^2^ = 0.49, and F = 5.01, df = 5, P < 0.0001, R^2^ = 0.38, for males and females, respectively, Fig. 3 A-B).

### 3.4 Allocation of time to feeding

All the experimental flies spent more time foraging on Full diets (Fig. 4 A-B). However, the longest time spent on Full diets was recorded in aposymbiotic flies regardless of drop size. Overall, aposymbiotic females spent on average 46.27±2.15 minutes on Full diets compared to 28.43±2.49 minutes by symbiotic females (F = 94.52, df = 5, P = 0.023, R^2^ = 0.96, Fig 4 A-B). Similarly, aposymbiotic males spent 38.27±4.15 minutes on Full diets, compared to 19.43±2.49 minutes for symbiotic males (F = 33.14, df = 5, P = 0.041, R^2^ = 0. 92, Fig. 4 A-B).

**FIG. 4.**
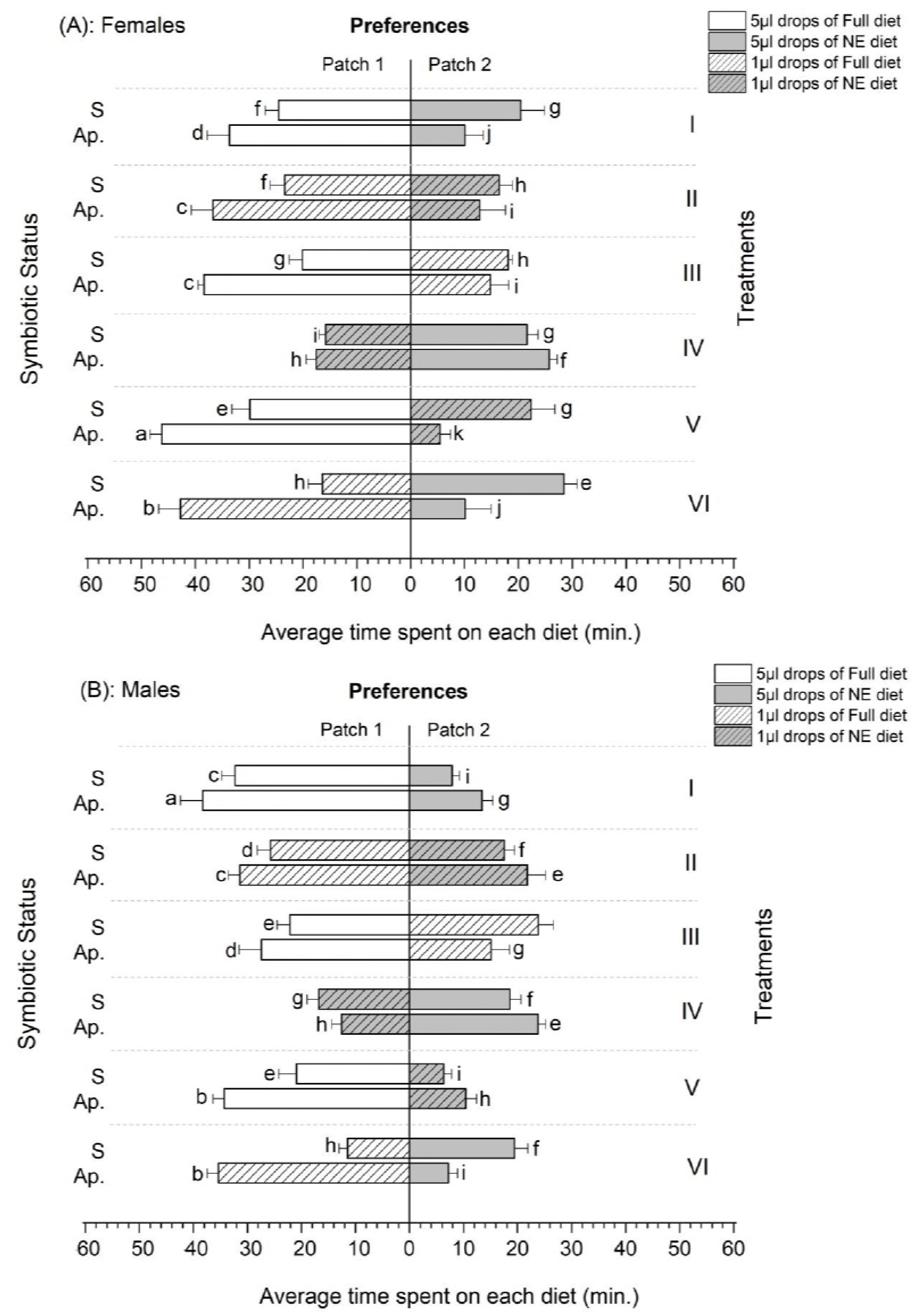
Feeding duration of *Bactrocera dorsalis* females (A) and males (B) under different foraging environments (Treatments). S: Symbiotic and Ap.: Aposymbiotic. Mean bars with different letters within and between treatments are statistically different after parametric Duncan’s Multiple Range Range Test at P = 0.05.

A comparison based on drop size also revealed a significantly longer time spent on high volume drops by aposymbiotic flies, except in treatment VI (Fig. 4 A-B). Aposymbiotic flies feeding on NE diets recorded the shortest time spent, with an average time of 5.43±1.95 minutes in females compared to 46.27±2.15 minutes on Full diets and 7.9± 1.33 minutes in males compared to 38.27±4.15 minutes (Fig. 4 A-B).

## 4. Discussion

Foraging entails decision making, whereby each individual must consider trade-offs between energetic and nutrient gain, and the time and risk associated with this activity (2). Furthermore, when organisms need to ingest nutrients from various food sources, behavioral mechanisms that optimize intake come into play (4). Gut bacteria have been implicated in this decision making process, both in invertebrates (20, 22) and vertebrates (12, 41).

In our experiments, suppressing the microbiome resulted in significant changes of the foraging behavior of both male and female flies. Aposymbiotic flies responded faster to the diets offered in experimental arenas, spent more time feeding, ingested more drops of food, and were constrained to feed on time consuming patches (containing small drops of food), when these offered the full complement of amino acids (treatment VI)).

These findings join a number of recent studies in elaborating the effect of gut bacteria on different stages of the nexus linking the gut to behavior, and significantly, extend this nexus to patterns of active foraging.

Aposymbiotic flies responded at a higher rate and with greater celerity to the experimental foraging arenas, compared to symbiotic flies (Figure 2, A & B). This suggests that the absence of bacteria in the gut affected the motivational state of these flies, by lowering the response threshold to visual and olfactory stimuli associated with food. In insects, response thresholds to external chemical and visual stimuli are modulated by physiological status (6), which in turn is affected by the presence and composition of the gut microbiome (15, 16). Our results join previous studies in showing how the presence or absence of intestinal bacteria can affect behavioral thresholds (7, 20, 42).

The flies in our experiments, both symbiotic and aposymbiotic, were maintained on a Nitrogen free diet prior to their introduction into the foraging arenas. Previous work on tephritids has established that the gut microbiome is capable of transforming non-essential amino acids (and other intractable sources of Nitrogen), into the building blocks necessary for reproduction and development (32, 33) (MA and CYN, unpublished data). Accordingly, we hypothesized that, when presented with a choice between a diet containing only the non-essential amino acids and a diet containing all amino acids, symbiotic flies would behave in a manner consistent with optimal foraging theory, while aposymbiotic flies would be constrained to forage preferentially on Full diets, at the cost of significantly extending the amount of time spent foraging. Indeed, the results of our experiment support these predictions. Symbiotic flies significantly spent less time feeding than aposymbiotic flies, and achieved this (energetic and risk averse) saving by feeding on large drops, irrespective of the diet they contained. Conversely, aposymbiotic flies spent more time ingesting food drops, and were compelled to seek drops containing the full diet, even when this choice entailed ingesting a large number of small drops when large (but essential amino acid deficient) drops were available, as in treatment VI (Figures 3 & 4).

We suggest that the next dimension to be explored in this context is a life history one. In monophagous species obligatory gut symbionts enable exploitation of otherwise toxic hosts during the larval stage (28, 43), or facultatively enable expansion of the native host range (44). Empirically, the microbiome of polyphagous tephritids is more varied than that of monophagous species (23, 28, 45).

Thus the ability of the microbiome to contribute to the larval phase may come at a cost during the adult phase, when a more varied microbiome may be more advantageous. In this study we examined the foraging behavior of adult oriental fruit flies, a polyphagous species, with a varied microbiome (24, 38, 39). In future studies we will examine the performance of monophagous flies in similar experimental foraging environments.

## 5. Conclusion

The results of our study support the emergent paradigm of the effect of gut bacteria on their hosts, which extends from basic metabolism to the nervous system, affecting gustatory thresholds, feeding behavior and ultimately (as shown here), to patterns of foraging in imbalanced nutritional environments. In future studies we plan to add a life history dimension to these observations.

## Data accessibility

This manuscript has no additional data.

## Author’s contribution

CYN, BY and EJ conceived and designed the study. MA, AAA and JN conducted the experiments and analyzed the data. KK and CZZ helped in statistical analysis. BY and CYN edited and revised the manuscript. All authors read and approved this submission.

